# A high-quality functional genome assembly of *Delia radicum* L. (Diptera: Anthomiidae) annotated from egg to adult

**DOI:** 10.1101/2021.06.11.447147

**Authors:** Rebekka Sontowski, Yvonne Poeschl, Yu Okamura, Heiko Vogel, Cervin Guyomar, Anne-Marie Cortesero, Nicole M. van Dam

## Abstract

Belowground herbivores are overseen and underestimated, even though they can cause significant economic losses in agriculture. The cabbage root fly *Delia radicum* (Anthomyiidae) is a common pest in *Brassica* species, including agriculturally important crops, such as oil seed rape. The damage is caused by the larvae, which feed specifically on the taproots of *Brassica* plants until they pupate. The adults are aboveground-living generalists feeding on pollen and nectar. Female flies are attracted by chemical cues in *Brassica* plants for oviposition. An assembled and annotated genome can elucidate which genetic mechanisms underlie the adaptation of *D. radicum* to its host plants and their specific chemical defenses, in particular isothiocyanates. Therefore, we assembled, annotated and analyzed the *D. radicum* genome using a combination of different Next Generation Sequencing and bioinformatic approaches. We assembled a chromosome-level *D. radicum* genome using PacBio and Hi-C Illumina sequence data. Combining Canu and 3D-DNA genome assembler, we constructed a 1.3 Gbp genome with an N50 of 242 Mbp and 6 pseudo-chromosomes. To annotate the assembled *D. radicum* genome, we combined homology-, transcriptome- and *ab initio*-prediction approaches. In total, we annotated 13,618 genes that were predicted by at least two approaches. We analyzed egg, larval, pupal and adult transcriptomes in relation to life-stage specific molecular functions. This high-quality annotated genome of *D. radicum* is a first step to understanding the genetic mechanisms underlying host plant adaptation. As such, it will be an important resource to find novel and sustainable approaches to reduce crop losses to these pests.

## 1. INTRODUCTION

The cabbage root fly, *Delia radicum* L. (Diptera; Anthomyiidae), is a severe pest in agriculture. The family Anthomyiidae, or flower flies, is a large family mainly occurring in the northern hemisphere. Adult *D. radicum* flies live aboveground and feed on nectar (Figure 1, (Gouinguene & Städler, 2005; Peter Roessingh & Städler, 1990)). The females oviposit next to or on the root crown of brassicaceous plants. After the eggs have hatched, the larvae occupy a new habitat and move into the soil to mine into the tap roots (Figure 1). After passing through three instars in about 20 days, the larvae move back to the soil to pupate (Capinera, 2008).

**Figure 1.**
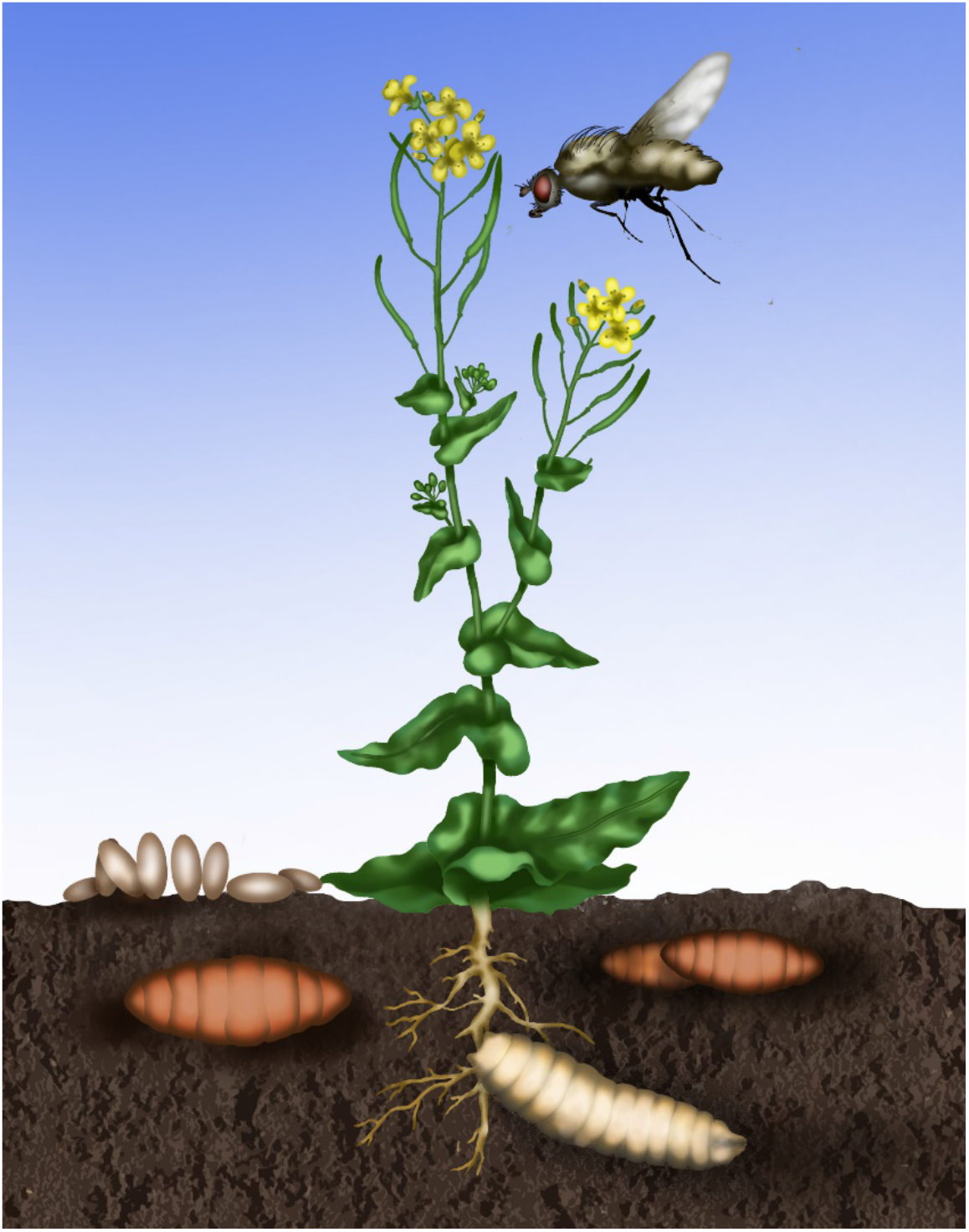
Schematic illustration of the life stages and their habitats of the cabbage root fly *Delia radicum*. Adult flies are attracted by their host plants for food consumption and oviposition. Eggs are deposited on the soil, where the larvae hatch. Larvae dig into the soil to feed on the roots until they pupate. After completing metamorphosis, adult flies make their way above ground to feed on pollen and nectar, and to reproduce. Picture: Jennifer Gabriel

As its common name “cabbage root fly” already indicates, *D. radicum* is a specialized herbivore on Brassicaceae, the cabbage and mustard family. This plant family contains several agriculturally important crops, such as broccoli, turnip, Pak Choi and rapeseed. Although they are specialists on Brassicaceae, females prefer some plant species of this family more for oviposition than others (Lamy et al., 2018). The female flies are attracted to the plant by specific volatile organic compounds, such as sulfides and terpenes (Ferry et al., 2007; Kergunteuil, Dugravot, Danner, van Dam, & Cortesero, 2015). Upon contacting the plants, the females decide to oviposit based on chemical cues, in particular the presence of glucosinolates (Gouinguené & Städler, 2006). The larvae are well adapted to deal with the glucosinolate-myrosinase defense system that is specific to the Brassicaceae (Hopkins, van Dam, & van Loon, 2009). Glucosinolates are sulfur-containing glycosylated compounds, which are stored in the vacuoles of cells localized between the endodermis and phloem cells (Kissen, Rossiter, & Bones, 2009). The roots of Brassica species contain high levels of glucosinolates, in particular 2-phenylethylglucosinolate (van Dam, Tytgat, & Kirkegaard, 2009). Glucosinolates can be converted by the enzyme myrosinase into pungent and toxic products, such as isothiocyanates (ITCs) and nitriles which deter generalist herbivores (Kissen et al., 2009). The myrosinase enzymes are stored in so called myrosin cells (Kissen et al., 2009). Upon tissue damage, either by mechanical damage or by herbivores, such as *D. radicum* larvae, the glucosinolates and myrosinases mix. This results in the formation of various conversion products, including ITCs, nitriles and sulfides (Crespo et al., 2012; Danner et al., 2015; Wittstock & Gershenzon, 2002).

Indeed, *D. radicum* larvae can successfully infest the roots of a wide range of Brassicaceae (Finch & Ackley, 1977; Tsunoda, Krosse, & van Dam, 2017). The damage the feeding larvae cause leads to substantial fitness loss in wild plants and yield reduction in crops. In rapeseed, *D. radicum* infestation reduces seed numbers and seed weight (Griffiths, 1991; McDonald & Sears, 1992). The annual economic losses due to *D. radium* infestation in Western Europe and Northern America are estimated to be 100 million $ (Wang, Voorrips, Steenhuis-Broers, Vosman, & van Loon, 2016).

Controlling *D. radicum* in agriculture is a major challenge. Natural resistance to this specialist herbivore has not been identified in currently used cultivars yet (Ekuere et al., 2005) and several effective synthetic insecticides, such as neonicotinoids, have been banned from use due to environmental concerns (Allema B, Hoogendoorn M, van Beek J, & P, 2017). Moreover, pesticide resistance has already developed in this species, for example against chlorpyrifos (van Herk et al., 2016). Alternative and more sustainable pest management strategies are urgently needed. Heritable natural resistance to *D. radicum* is present in wild brassicaceous species, but introgression of these traits may be hampered by crossing barriers and linkage of resistance with undesired traits (Ekuere et al., 2005; Wang et al., 2016). Several studies examined the application of entomopathogenic fungi, natural predators or parasitoids, mixed cropping and soil microbes to better control *D. radicum* (Bruck, Snelling, Dreves, & Jaronski, 2005; Dixon, Coady, Larson, & Spaner, 2004; Fournet, Stapel, Kacem, Nenon, & Brunel, 2000; Kapranas, Sbaiti, Degen, & Turlings, 2020; Lachaise et al., 2017; Neveu, Krespi, Kacem, & Nénon, 2000). Even though each of these measures may reduce *D. radicum* infestations, they cannot prevent yield loss as effectively as synthetic pesticides.

To understand the interaction of *D. radicum* with its host plants, the chemical ecology of this plant-herbivore interaction has been intensively studied over the last decades. These studies analyzed aspects ranging from the chemosensory mechanisms of host plant attraction and oviposition choice to herbivore-induced plant responses and interactions with predators and parasitoids (Ferry et al., 2007; Gouinguene & Städler, 2005; Hopkins et al., 2009; Kergunteuil et al., 2015; P Roessingh et al., 1992). However, the genetic mechanisms underpinning host-plant adaptation of *D. radicum* are unknown. An accurate and well-annotated genome can reveal genetic mechanisms underpinning adaptation of *D. radicum* to its host’s chemical defenses. Especially, understanding the preference of the different agricultural relevant life stages (adults and larvae) which occur in separate habitats (above- and belowground) on the genetic level expands our understanding of herbivore-plant interactions. These mechanisms can also be an important starting point to develop novel approaches, such as species-specific dsRNA-based pest control strategies. So far, a genome of this species has not been published.

Here, we assembled and annotated a *de novo*, chromosome-level scaffolded genome of *D. radicum* using PacBio and Hi-C Illumina sequencing. We used three different approaches to annotate the genome; Cufflinks, which uses transcript assembly; GeMoMa, which is homology-based, and BRAKER, for additional prediction of genes not covered by the first two methods. Generated RNASeq data of all four life stages (eggs, larvae, pupae, adults) and two relevant stress factors (heat stress in adults, plant toxin stress in larvae), allowed us to validate predicted genes and to identify specific gene families which were expressed in each of the life stages.

## 2. MATERIALS AND METHODS

### 2.1 Sample material

A starting culture of *D. radicum* was provided by Anne-Marie Cortesero, University of Rennes, France in 2014. It originated from pupae collected in a cabbage field in Brittany, France, (48°6′31″ N, 1°47′1″ W) the same year. More than five thousand pupae were collected to start the original culture and ~50 individuals from this culture were sent to iDiv. A permanent culture was established in our lab under constant conditions (20 ± 2°C, 85 ± 10 % RH, 16L:8D) in a controlled environment cabinet (Percival Scientific, Perry, Iowa, USA) resulting in an inbreeding line of over 60 generations. Adult flies were reared in a net cage and fed with a 1:1 milk powder-yeast mixture and a water-honey solution, which was changed three times a week. Water was provided ad libitum. Eggs were placed in a 10×10×10 cm plastic box filled with 2 cm moistened, autoclaved sand and a piece of turnip. Once the larvae hatched, old turnip pieces were removed and exchanged with new turnip every other day and the sand was moistened when necessary. After the third instar, the larvae crawled into the sand, where they pupated. The pupae were collected by flooding the box with water, collecting the floating pupae and placing them into the adult fly cage until eclosion.

Species identification was performed using a fragment of the cytochrome oxidase I (COI) gene as a molecular marker generated with the universal COI primer pair HCO and LCO (Folmer et al. 1994). The sequence was submitted to BLAST online using the BLASTn algorithm (Retrieved from https://blast.ncbi.nlm.nih.gov/Blast.cg). The top three hits matched with *D. radicum* COI accessions (MG115888.1, HQ581775.1, GU806605.1) with an identity of more than 98.45 %.

### 2.2 Genome sequencing

#### 2.2.1 Sampling, DNA extraction and PacBio sequencing

For PacBio sequencing, 18 randomly collected, fully matured *D. radicum* adults were frozen and stored at −80°C. To sterilize the surface, the flies were incubated for 2 min in bleach (2 %), transferred to sodium thiosulfate (0.1 N) for neutralization, washed three times in 70 % ethanol and once in autoclaved dd water. To reduce contamination by microorganisms from the gut, we extracted total DNA from the head and the thorax of the adults, using a phenol-chloroform extraction method according to the protocol of the sequencing facility (Figure S1). We pooled three individuals per extraction and checked the DNA quality using gel electrophoresis (0.7 % agarose gel). DNA purity was assessed using a NanoPhotometer® P330 (Implen, Munich/Germany) and DNA quantity using a Qubit dsDNA BR assay kit in combination with a Qubit 2.0 Fluorometer (Invitrogen, Carlsbad, CA/USA). The DNA of all samples was pooled for the sequencing library. Library preparation and sequencing was provided by the facility of the Max Planck Institute of Molecular Cell Biology and Genetics, Dresden/Germany on a PacBio Sequel. A total of 16 SMRT cells were processed and 6,539,960 reads generated. Due to the pooling of several females and males, we expected the reads to be highly heterozygous, which we considered for the assembly process.

#### 2.2.2 Sampling, DNA extraction and Hi-C Illumina sequencing

For Hi-C Illumina sequencing fresh *D. radicum* pupae from the above culture were randomly selected. A total of 10 pupae were chopped into small pieces with a razor and resuspended in 3 ml of PBS with 1% formaldehyde. The homogenized sample was incubated at RT for 20 min with periodic mixing. Glycine was added to the sample buffer to 125 mM final concentration and incubated at RT for ~15 min with periodic mixing. The homogenized tissue was spun down (1000 × g for 1 min), rinsed twice with PBS, and pelleted (1000 × g for 2 min). After removal of the supernatant, the tissue was homogenized to a fine powder in a liquid nitrogen-chilled mortar with a chilled pestle. Further sample processing and sequencing was performed by Phase Genomics (Seattle, WA, USA) on an Illumina HiSeq 4000, generating a total of 181,752,938 paired end reads (2 × 150 bp).

### 2.3 Genome size estimation

A karyotyping study determined that *D. radicum* is a diploid organism with 2n = 12 chromosomes (Hartman & Southern, 1995). To obtain a reliable estimate of the *D. radicum* genome size, we used flow cytometry based on a method using propidium iodide-stained nuclei (Spencer Johnston Lab, Texas A&M, USA; (Hare & Johnston, 2011). The haploid genome sizes were estimated to be 1239.0 ± 27.5 Mbp for females (N = 4) and 1218.0 ± 4.0 Mbp for males (N = 50).

### 2.4 Genome assembly and completeness

#### 2.4.1 PacBio data processing

Raw PacBio reads in bam file format were converted into fasta files by using samtools (version 1.3.1) (Li et al., 2009) as part of the SMRT link software (version 5.1.0, https://www.pacb.com/support/software-downloads/). Extracted raw PacBio reads (6,539,960 reads) were checked for potential contaminations with prokaryotic DNA by applying EukRep (version 0.6.2) (West, Probst, Grigoriev, Thomas, & Banfield, 2018) with default parameter settings. Only reads classified as eukaryotic (4,454,601 reads) were used for the *de novo* genome assembly.

#### 2.4.2 *De novo* genome assembly

The long-read assembler Canu (version 1.9) (Koren et al., 2017) was used to generate a *de novo* genome assembly from contamination-free PacBio reads. The Canu pipeline, including read error correction and assembly, was started with setting parameters based on the estimated genome size (genomeSize=1200m) and the use of not too short (minReadLength=5000) and high quality (stopOnReadQuality=true) PacBio reads, addressing the overlapping of sequences (minOverlapLength=1000 corOutCoverage=200), and accounting for the expected high heterozygosity rate of the *D. radicum* genome (batOptions=-dg 3 -db 3 -dr 1 -ca 500 -cp 50). The latter parameters were selected to prevent the haplotypes from being collapsed during the assembly process.

#### 2.4.3 Polishing and purging

To improve the sequence quality of the raw genome assembly, we performed two rounds of polishing. All eukaryotic raw PacBio reads were aligned with pbalign (version 0.3.1 and default parameter settings) and these results were used for sequence polishing with Arrow (version 2.2.2 and default parameter settings). Both programs are part of the SMRT link software (version 5.1.0, https://www.pacb.com/support/software-downloads/). To detect and remove duplications in the assembled contigs, we applied purge_dups (version 1.2.3) (Guan et al., 2020) on the polished assembly. We ran the first three steps of the purge_dups pipeline with default parameters and the last step with additional setting “-e -c” to allow only clipping at the end of contigs and retaining high coverage contigs.

#### 2.4.4 Chromosome-scale scaffolding

Hi-C Illumina reads were aligned to the purged assembly with the Juicer pipeline incorporating juicer_tools (version 1.22.01) (Durand et al., 2016), “-s DpnII” and a restriction site file (generated with the generate_site_positions.py script contained in juicer) provided by “-z” option. The sequences of the purged assembly were scaffolded with the Juicer output on Hi-C read alignments into chromosome-scale super-scaffolds applying the 3D-DNA genome assembler (version 18011) (Dudchenko et al., 2017) with the additional setting of “--splitter-coarse-stringency 30 --gap-size 100”. This resulted in the final genome assembly from *D. radicum*.

#### 2.4.5 Evaluating genome completeness

We used BUSCO v4 (4.0.5) (Seppey, Manni, & Zdobnov, 2019) to analyze the completeness of the final and intermediate genome assemblies. Three different gene sets, insecta_odb10.2019-11-20, endopterygota_odb10.2019-11-20, and diptera_odb10.2019-11-20, representing different levels in evolutionary relatedness were considered in the evaluation process. These three gene sets comprise 1367, 2124, or 3285 orthologous genes, respectively.

#### 2.4.6 Exclusion of Hi-C scaffolds

While assembling the *D. radicum* genome we co-assembled the complete genome of *Wolbachia* (Hi-C scaffold 7) a common endosymbiont in arthropods. To obtain a contamination-free final assembly, we excluded Hi-C scaffolds 7, 146 and 370 and trimmed Hi-C scaffold 6 after position 12,881,041 that were annotated to be contaminated with *Wolbachia* sequences during the NCBI validation process.

### 2.5 Phylogeny - Comparative genomics based on BUSCOs

Phylogenetic analyses were done with BuscoOrthoPhylo (https://github.com/PlantDr430/BuscoOrthoPhylo) which is a wrapper script to concatenate and align protein sequences, and to construct a phylogenetic tree based on single-copy BUSCO genes. BUSCOs of the endopterygota_odb10.2019-11-20 gene set, consisting of 2124 genes, were used as the basis for the analysis. In the initial phase complete single-copy BUSCO genes which were shared by 10 selected species (Table 1), were computed. Protein sequences of the shared genes were extracted and concatenated for each species. MAFFT aligner (version 7.475) (Katoh, Misawa, Kuma, & Miyata, 2002) was run on concatenated FASTA file(s) and finally RAxML (version 8.2.12) (Stamatakis, 2014) with “-rx_p_sub PROTGAMMAWAG” as model and “-b 100” bootstrap steps was used to reconstruct the phylogenetic tree. The resulting findings were visualized in a phylogenetic tree using Phylo.io (Robinson, Dylus, & Dessimoz, 2016).

**Table 1.** 9 selected insect species. 9 insect species (4 Diptera, 4 Lepidoptera, and 1 Coleoptera species) selected for comparative genomics and phylogenetic analyses. Insect species were chosen according to their phylogenetic relatedness to *D. radicum*, or because they shared the same host plant range with *D. radicum* or because they are also common pests in agriculture. All 9 species are fully sequenced and annotated, and information can be obtained from National Center for Biotechnology (https://www.ncbi.nlm.nih.gov).

| Order | Species | NCBI taxid | Names | RefSeq ID | Reason for selection |
| --- | --- | --- | --- | --- | --- |
| Diptera | *Anopheles gambiae* | 180454 | African malaria mosquito | GCF_000005575.2 | phylogenetically related |
| Diptera | *Drosophila melanogaster* | 7227 | fruit fly | GCF_000001215.4 | phylogenetically related |
| Diptera | *Lucilia cuprina* | 7375 | Australian sheep blowfly | GCF_000699065.1 | phylogenetically related |
| Diptera | *Musca domestica* | 7370 | house fly | GCF_000371365.1 | phylogenetically related |
| Lepidoptera | *Manduca sexta* | 7130 | tobacco hornworm | GCF_000262585.1 | common pests on crop plants |
| Lepidoptera | *Pieris rapae* | 64459 | cabbage white | GCF_001856805.1 | sharing host plant |
| Lepidoptera | *Plutella xylostella* | 51655 | diamondback moth | GCF_000330985.1 | sharing host plant |
| Lepidoptera | *Spodoptera litura* | 69820 | Tobacco cutworm | GCF_002706865.1 | common pests on crop plants |
| Coleoptera | *Tribolium castaneum* | 7070 | red flour beetle | GCF_000002335.3 | common pests on stored grains |

### 2.6 Sampling, RNA extraction and Transcriptome sequencing

All life stage samples were collected from the laboratory culture (section 2.1). We used three replicates per life stage and condition. For the egg stage, we collected 25 mg eggs (laid within 24 h) per replicate. For the larval stage, we collected 18 randomly selected second instar larvae. Nine of the selected larvae were fed on a semi-artificial diet, containing yeast, milk powder, freeze dried turnip, agar (2:2:2:1) and 90% water. The other nine larvae were reared on the same diet containing 0.4 mg phenylethyl isothiocyanate/g diet. All larvae received freshly prepared diet every other day. After 7 days, larvae were shock-frozen at −80°C and pooled into batches containing 3 larvae forming three biological replicates per treatment. For the pupal stage, we randomly selected nine freshly formed pupae and pooled them into three biological replicates of three pupae each. For the adult stage, we collected 18 fully developed random adults. Nine individuals were exposed to 35°C (Michaud, Marin, Westwood, & Tanguay, 1997) for 2 hours, whereas the control adults were kept under normal conditions. We pooled three adults for one replicate, resulting in three replicates for control and elevated temperature treatment.

All samples were surface sterilized using the same procedure as described for the adult flies. We extracted the total RNA of the larval stage using ReliaPrep RNA Tissue Miniprep kit (Promega, Madison, WI/USA) according to the supplier’s recommended protocol. Total RNA of all further samples was extracted using TRIzol (Life technologies, Carlsbad, CA/USA) according to the supplier’s recommended protocol. Qualitative and quantitative RNA assessment of all samples was done by gel electrophoresis (1% agarose), a NanoPhotometer® P330 (Implen, Munich/Germany) and a Qubit 2.0 (Invitrogen, Carlsbad, CA/USA).

Library preparation and sequencing of the larval samples (control and stressed larvae) were performed by the Deep Sequencing group of Biotech TU Dresden/Germany on an Illumina NextSeq next generation sequencer. The poly(A) enriched strand specific libraries generated for all samples ran on one lane generating in total 400 Mio of 75 bp paired end reads. Egg, pupal and adult (control and stressed) samples were sequenced by Novogene (Hong Kong/China) with strand specific library preparations and sequencing on an Illumina NovaSeq 6000 next generation sequencer, generating 20 Mio paired-end (2 × 150 bp) reads per sample.

### 2.7 Genome annotation – prediction of protein-coding genes

#### 2.7.1 Mapping of Transcriptome data

Including RNASeq data can improve the quality of gene predictions as optional input by several gene prediction algorithms. We mapped the *D. radicum* RNASeq data of the 18 samples, consisting of six conditions (4 life stages and 2 stress treatments) with three replicates each to the *D. radicum* genome with STAR (version 020201) (Dobin et al., 2012) and store mapping results in bam files.

#### 2.7.2 Homology-based gene prediction

Homology-based GeMoMa (version 1.6.4 und 1.7.2) (Keilwagen, Hartung, Paulini, Twardziok, & Grau, 2018; Keilwagen et al., 2016) gene predictions the *D. radicum* genome were performed based on the annotated genomes of four Diptera species (*Anopheles gambiae*, *Drosophila melanogaster*, *Lucilia cuprina*, and *Musca domestica*), four Lepidoptera species (*Manduca sexta*, *Pieris rapae*, *Plutella xylostella*, and *Spodoptera litura*), and one Coleoptera species (*Tribolium castaneum*) obtained from NCBI (Table 1). For each of these nine species, extracted CDS were aligned with MMseqs2 (version 11.e1a1c) (Steinegger & Söding, 2017) to the *D. radicum* genome sequence with parameter values suggested by GeMoMa. Alignments and RNASeq mappings were used for predictions of gene models in the genome with GeMoMa and default parameters, separately for each species and by incorporating mapped RNASeq data for refining intron boundaries. The resulting nine gene annotation sets were filtered and merged using the GeMoMaAnnotationFilter (GAF) with “f=”start==‘M’ and stop==‘*’” atf=‘’”. Only transcripts of genes starting with the start codon “M(ethionine)” and ending with a stop codon “*” were considered and all isoforms were retained. We finally predicted and added UTR annotations to the resulting filtered set of transcripts by using the GeneAnnotationFinalizer with “u=YES rename=NO”, which is also part of the GeMoMa suite.

#### 2.7.3 Transcriptome assembly - RNA-Seq-based gene predictions

To assemble one transcriptome per life stage and condition, we merged the read mappings (bam files) of the three replicates per condition and life stage. For the transcriptome assembly of the mapped RNASeq data, we used Cufflinks (version 2.2.1) (Trapnell et al., 2010). Initially, soft-clipped read mappings were clipped, and assembled to six transcriptomes using Cufflinks with default parameters and “-fr-firstrand”. The resulting six transcriptomes were subjected to Cuffmerge, which is part of the Cufflinks toolbox, to generate a single master transcriptome. While Cufflinks assembled transcripts with exon annotation, missing coding regions and UTRs were identified with TransDecoder (version 5.5.0, https://github.com/TransDecoder/TransDecoder). Finally, predicted transcripts were filtered for a proper start and end of protein coding transcripts and retaining the UTR annotations by applying the GAF with parameters “f=”start==‘M’ and stop==‘*’” atf=’’ aat=true tf=true”.

#### 2.7.4 *Ab initio* gene prediction

Additionally, we aimed to predict genes not covered by the homology-based and the transcriptome-based approach, due to a lack of homology or because of no or low expression under the specific conditions of the sampled life stages. To obtain such *ab initio* gene predictions, we ran RepeatMasker (version 4.1.0, http://www.repeatmasker.org) with RMBlast (version 2.10.0, http://www.repeatmasker.org/RMBlast.html) and “-species insecta -gff -xsmall” to find and mask repetitive sequences annotated for insects in the RepeatMasker repeat database. For *ab initio* prediction of protein-coding genes on the masked genome sequences, we ran BRAKER (version 1.9) (Brůna, Hoff, Lomsadze, Stanke, & Borodovsky, 2021; Katharina J. Hoff, Lange, Lomsadze, Borodovsky, & Stanke, 2015; Katharina J Hoff, Lomsadze, Borodovsky, & Stanke, 2019), which combines GeneMark (version 4.59_lic) (Lomsadze, Burns, & Borodovsky, 2014) and AUGUSTUS (version 3.4.0) (Stanke, Schöffmann, Morgenstern, & Waack, 2006) with “--gff3--softmasking” and provided the mapped RNASeq data as hints for initial training of gene models and gene predictions.

Predicted transcripts were filtered for proper start and end of protein coding transcripts by applying the GAF with “f=”start==‘M’ and stop==‘*’” atf=‘’”. Finally, UTR annotations were predicted and added using GeneAnnotationFinalizer with “u=YES rename=NO”.

#### 2.7.5 Final genome annotation and completeness evaluation

We ran GeMoMa`s GeMoMaAnnotationFilter (GAF) with “f=‘’ atf=‘’ tr=true aat=true” to integrate the predicted gene models from all three applied approaches, the homology-based, the RNASeq-based and *ab initio* gene prediction approach, and yield a master gene annotation file for the *D. radicum* genome. As gene-related features we include mRNA, CDS, five_prime_UTR, and three_prime_UTR specificities in the annotation file and several attributes that gave additional information on the predicted transcripts and can be used for user-specific filtering.

We evaluated the completeness of the final set of protein-coding genes with BUSCO v4 as we did for the evaluation of the completeness of the genome (section 2.4.5), but set “-m proteins”.

### 2.8 Functional annotation

Predicted *D. radicum* protein sequences were subjected to PANNZER2 (Protein ANNotation with Z-scoRE) (Törönen, Medlar, & Holm, 2018), which predicts functional descriptions and GO classes. Additionally, extracted protein sequences were subjected to InterProScan (version 5.45-80.0.) (Blum et al., 2020; Jones et al., 2014) and scanned for information on protein family and domains in all member data bases (-appl CDD, HAMAP, PANTHER, Pfam, PIRSF, PRINTS, ProDom, PROSITEPATTERNS, SMART, TIGRFAM, Gene3D, SFLD, SUPERFAMILY, MobiDBLite) and for GO- or pathway annotation (“-goterms-iprlookup -pa”).

GO terms annotated for transcripts with PANNZER2 and InterProScan were merged. Additionally, to get a functional annotation per gene, we merged the annotations of all respective transcripts.

### 2.9 Synteny analysis

Annotated CDSs of *D. melanogaster* (Table 1) were extracted and aligned to the *D. radicum* genome with MMseqs2 (version 11.e1a1c) (Steinegger & Söding, 2017). Alignments were used for homology-based predictions of gene models in the *D. radicum* genome with GeMoMa (version 1.6.4) (Keilwagen et al., 2018) with default parameters. Predicted gene models were filtered using the GeMoMaAnnotationFilter (GAF) with “f=”start==‘M’ and stop==‘*’” atf=‘’”. Finally, a table containing the relation and positions of the gene models was generated with SyntenyChecker, which is part of the GeMoMa toolbox. Syntenic relationships of *D. radicum* to *D. melanogester* were visualized using Circos (version 0.69-9) (Krzywinski et al., 2009).

### 2.10 Analysis of life cycle data

We extracted the sequences of all annotated transcripts and quantified their abundances with kallisto (version 0.46.1, (Bray, Pimentel, Melsted, & Pachter, 2016)) with “-b 100” bootstraps and “—rf-stranded”. The abundances were imported into the statistical framework R (version 3.6.2) (R Core Team, 2020) for further analyses using the R package tximport (1.14.2) (Soneson, Love, & Robinson, 2015). Using tximport transcript-level, estimates for abundances were summarized for further gene-level analyses.

We denoted a gene as *expressed* if it had a TPM (transcript per million) value ≥ 1 in at least one of the 18 transcriptome samples. We refer to this set of genes as the “data set of expressed genes”. We called a gene present in a life stage or condition if it occurred in at least one replicate. This aggregation resulted in a matrix with six columns (four life stages, two conditions). These six sets were analyzed for life stage and condition-specific gene expression and also for intersections with the R package UpSetR (1.4.0) (Gehlenborg, 2019).

We performed Gene Ontology (GO) analyses of pre-defined gene sets using R (version 4.0.4) with the latest version of the R package topGO (2.42.0) (Alexa & Rahnenfuhrer, 2020) with GO.db (3.12.1) (Carlson, Falcon, Pages, & Li, 2020). We used Fisher’s exact test to identify over-represented GO terms. Raw p-values were corrected for multiple testing using the method proposed by Benjamini and Yekutieli (Benjamini & Yekutieli, 2001) implemented in p.adjust contained in the basic R package stats. To get an indication of which processes were active, we aggregated single significant GO-terms (adjusted p value < 0.05) into self-assigned generic categories. Results were visualized using the R package pheatmap (1.0.12) (Kolde, 2019). For visualization of the results for generic categories, we computed the relative frequency of GO terms determined in a pre-defined gene set for a generic category. The relative frequency was calculated by the number of significant GO terms in a gene set divided by the total number of GO terms that were sorted into the appropriate generic category.

We defined six gene sets for life stage (eggs, larva, pupa, adults) and condition-specific GO analysis (ITC, heat stress). For the analysis of the whole life cycle we determined genes that were exclusively expressed in one of the four life stages under control conditions. As we have additional stress conditions in the larval and the adult life stage, we extended the defined gene sets for these two life stages by genes contained in the intersection of both conditions (control and stress) within these stages. For condition-specific GO analysis, we additionally determined the genes exclusively expressed in the stressed condition of the larval and adult stage, respectively. Again, we also extended the stress-specific genes sets by the respective intersection gene set.

We clustered samples and genes contained in the data of expressed genes using the R package umap (0.2.7.0) (Konopka, 2020). UMAP (uniform manifold approximation and projection) is a technique to reduce dimensions and bring similar data vectors, samples (columns) or genes (rows) in close proximity. In our analyses we projected the data vectors in both cases in a two-dimensional space and tested different values for the size of the neighborhood (n_neighbors) and the minimal distance (min_dist) between data points (either samples or genes).

## 3 RESULTS AND DISCUSSION

### 3.1 Genome assembly

PacBio reads classified as eukaryotic (4,454,601 reads) were used for a contamination-free assembly of the *D. radicum* genome with Canu. We expected a high heterozygosity rate due to the pooling of multiple *D. radicum* individuals. Setting Canu parameters accordingly to prevent haplotypes from being collapsed during the assembly process, resulted in a raw assembly with the length of approximately 2.538 Gbp, which was nearly twice the size of the expected genome, an N50 contig of approximately 205.3 Kbp and in total 29,244 contigs (Table 2). By evaluating the completeness of the raw genome assembly with BUSCO (using three different sets of orthologous genes at different levels of evolutionary relatedness), the *raw* assembly revealed a completeness of at least 95.7 % for the Diptera (Figure 2b, Table S1) and for more than 98 % for the Endopterygota gene set (Table S2). These results showed a high completeness of the raw assembly, but also the existence of a reasonable percentage of duplicated sequences.

**Table 2.**
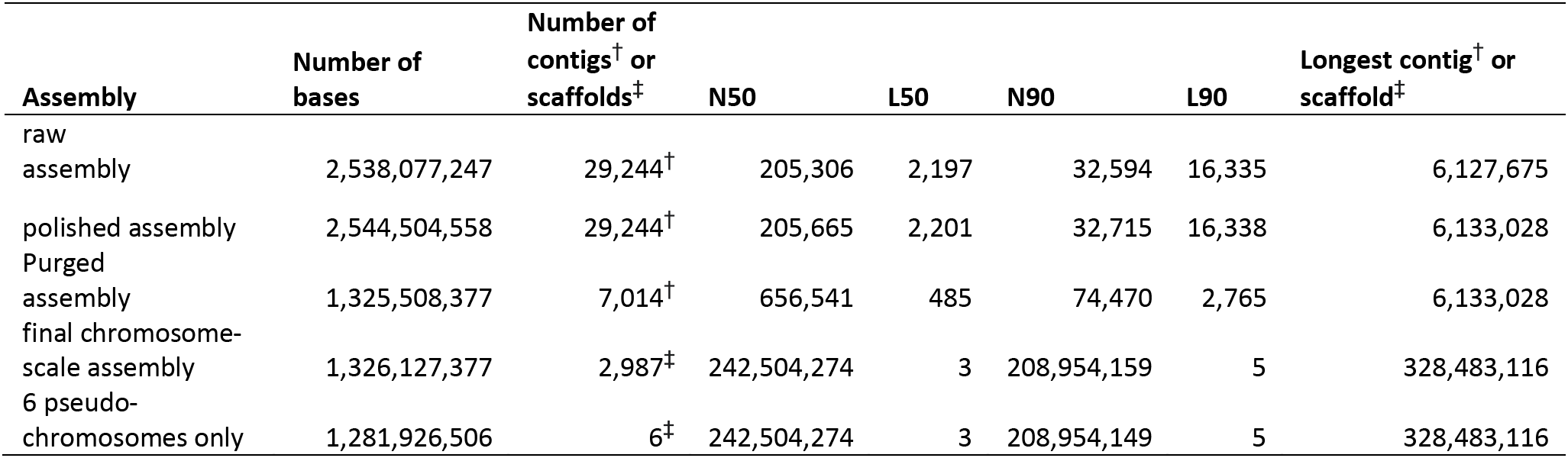
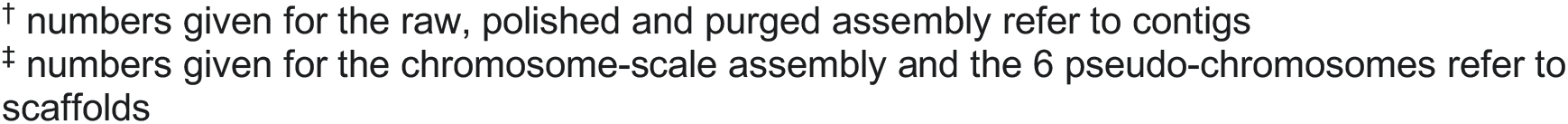
Summary of assembly statistics. The raw, polished, and purged assemblies are intermediate assemblies after PacBio read assembly with Canu, two rounds of polishing with Arrow, and purging with purge_dups. The final, chromosome-scale assembly, generated with the 3D-DNA genome assembly pipeline that assembled contigs of the purged assembly by integration of Hi-C Illumina reads into (chromosome-scale) scaffolds. The final chromosome-scale assembly contains 6,190 gaps of length 100 bp, whereby 6,188 gaps are located on the 6 pseudo-chromosomes.

| Assembly | Number of bases | Number of contigs<sup>†</sup> or scaffolds<sup>‡</sup> | N50 | L50 | N90 | L90 | Longest contig<sup>†</sup> or scaffold<sup>‡</sup> |
| --- | --- | --- | --- | --- | --- | --- | --- |
| raw assembly | 2,538,077,247 | 29,244<sup>†</sup> | 205,306 | 2,197 | 32,594 | 16,335 | 6,127,675 |
| polished assembly | 2,544,504,558 | 29,244<sup>†</sup> | 205,665 | 2,201 | 32,715 | 16,338 | 6,133,028 |
| Purged assembly | 1,325,508,377 | 7,014<sup>†</sup> | 656,541 | 485 | 74,470 | 2,765 | 6,133,028 |
| final chromosome-scale assembly | 1,326,127,377 | 2,987<sup>‡</sup> | 242,504,274 | 3 | 208,954,159 | 5 | 328,483,116 |
| 6 pseudo-chromosomes only | 1,281,926,506 | 6<sup>‡</sup> | 242,504,274 | 3 | 208,954,149 | 5 | 328,483,116 |

† numbers given for the raw, polished and purged assembly refer to contigs ‡ numbers given for the chromosome-scale assembly and the 6 pseudo-chromosomes refer to scaffolds

**Figure 2.**
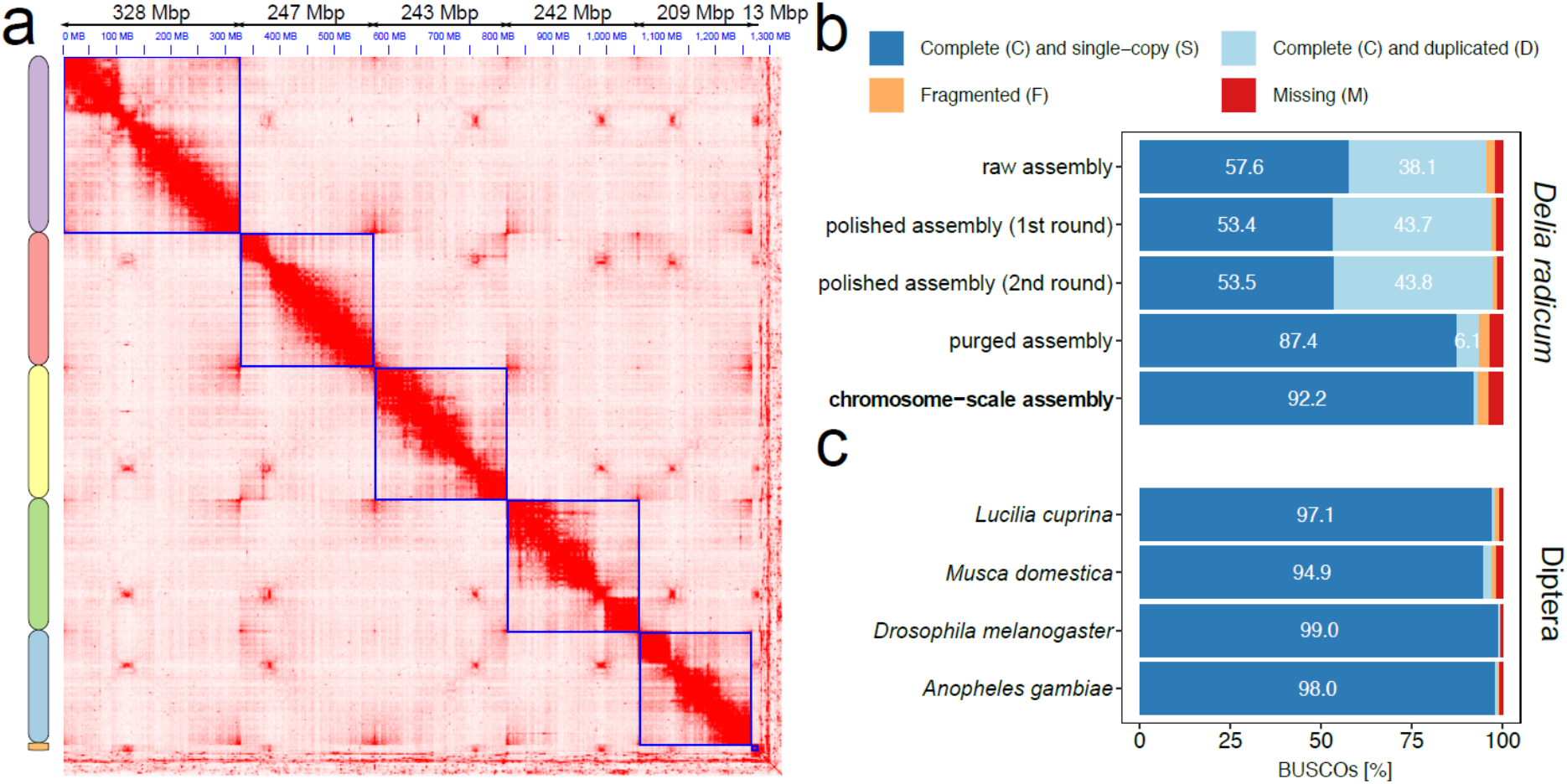
Chromosome-scale assembly of the *Delia radicum* genome. (a) Heatmap showing the Hi-C contacts map of the final chromosome-scale assembly, where the six chromosomes (six super-scaffolds) are indicated by the blue boxes. The chromosomes are ordered from largest to smallest; their concrete lengths are given in Mbp above the Hi-C map. (b) Barplot showing the result of BUSCO analyses of the intermediate and final assemblies using the ‘Diptera’ gene set containing 3,285 genes. Numbers in the bars give the percentage of genes found for the category indicated by the color of the bar. (c) Barplot showing the result of BUSCO analyses using the ‘Diptera’ gene set of four other dipteran species with published genomes. Numbers in the bars give the percentage of genes found for the category indicated by the color of the bar.

Improving the sequence quality of the raw genome assembly by performing two rounds of polishing with Arrow, increased not only the size of the assembly to approximately 2.544 Gbp (Table 2), but also the completeness of the *polished* assembly. Especially the percentage of complete genes in the BUSCO Diptera gene set increased to more than 97 %. Simultaneously, the number of duplicated genes increased (Figure 2b, Table S1).

Next, removing duplicated sequences in the polished assembly with purge_dups successfully reduced the size of the assembly to approximately 1.326 Gbp and a total of 7,014 contigs with an N50 of nearly 656.5 Kbp. The size of the *purged* assembly was already close to the experimentally determined genome size. By evaluating the completeness of the purged assembly, we observed a strong reduction in the percentage of duplicated genes, for the Diptera gene set to 6.1 % (Figure 2b, Table S1). As a side effect of removing sequences, the completeness of the gene sets dropped slightly to 93.5 % (Figure 2b, Table S1).

For the final *chromosome-scale* assembly, we scaffolded the contigs of the purged assembly with Hi-C data using Juicer and the 3D-DNA genome assembler. The resulting assembly comprised six chromosome-scale contigs (Figure 2a, Table S3), which was consistent with the number of chromosomes determined by karyotyping (Hartman & Southern, 1995), and 2,981 smaller, not-assembled contigs. The final assembly of the *D. radicum* genome yielded approximately 1.326 Gbp, where 96.67 % of the bases (nearly 1.281 Gbp) were anchored to the six pseudo-chromosomes. The size of the six pseudo-chromosomes ranged from one small chromosome with 13 Mbp to five larger chromosomes between 209 and 328.5 Mbp (Table S3). This is in line with the karyotype of *D. radicum*, which comprises five large and one much smaller chromosome (3.3 % of the large chromosomes’ size) (Hartman & Southern, 1995).

Validation of the final assembly with BUSCO (Table S1, S2) showed no considerable change in the number of complete genes, but the number of single-copy genes increased to 92.2 % (3,030 genes) while the number of duplicated genes decreased to 1.2 % (40 genes) for the Diptera gene set. The six pseudo-chromosomes along with the small contigs were used for all further analyses and are referred to as the *D. radicum* genome hereafter. The number of single-copy BUSCOs of the Diptera gene set in the *D. radicum* genome, was similar to those of other Diptera genomes (Figure 2c, Table S4), indicating that the chromosome-scale genome assembly of *D. radicum* was of comparable quality. Based on our findings, we can conclude that the final *D. radicum* chromosome-scale assembly was accurate, complete and without prokaryotic contamination.

### 3.2 Phylogeny and synteny

To examine the phylogenetic relationship of *D. radicum* to other insects, we compared complete single-copy BUSCOs of the Endopterygota gene set (comprising totally 2,124 genes) shared by the selected nine insect species belonging to Diptera (4), Coleoptera (1) and Lepidoptera (4, Table 1). We identified 1,217 (Table S5, Table S6) shared, and therefore conserved, single-copy genes (Figure 3a, Table S5). Reconstruction of the evolutionary relationships among these ten species based on the shared gene sets revealed that the root fly *D. radicum* was most closely related to the blow fly, *L. cuprina*, followed by the house fly, *M. domestica,* and the fruit fly, *D. melanogaster* (Figure 3a). These relations were consistent with their taxonomic position (Wiegmann et al., 2011).

**Figure 3.**
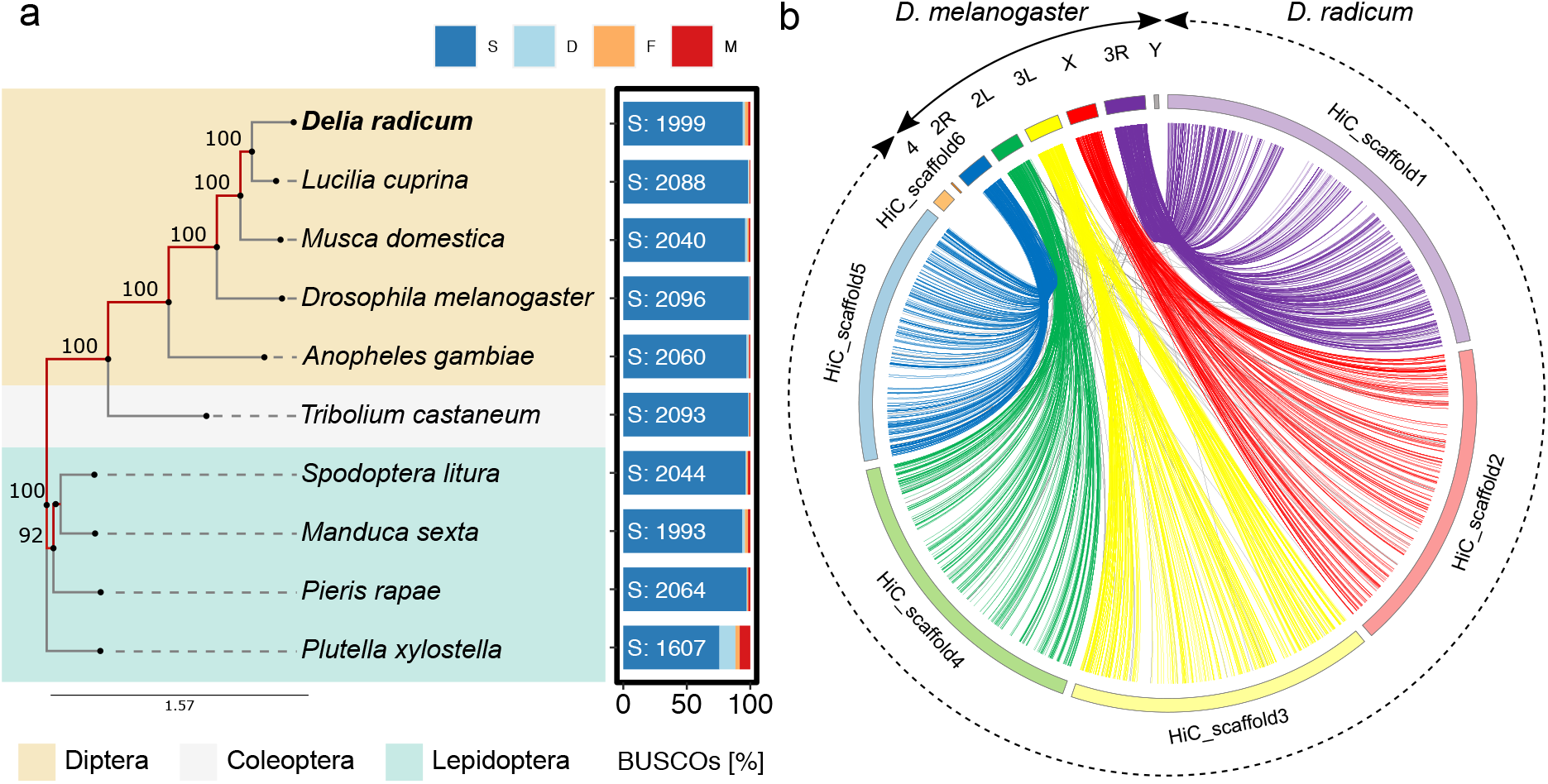
Phylogenetic analyses. (a) A phylogenetic tree reconstructed with RAxML on concatenated alignments of proteins of 1,271 genes of BUSCOs’ Endopterygota gene set (n=2,124) shared by all ten insect species. Tree reconstruction was done including 100 bootstrapping steps. The level of bootstrapping support is given at the edges. The barplot to the right of the phylogenetic tree shows BUSCO results of each species on the Endopterygota gene set, where S: number of complete single-copy BUSCO genes (dark blue bar); D: duplicated complete copy genes (light blue), F: fragmented genes (orange), M: missing genes (red). (b) A Circos plot linking genes on the assembled scaffolds of *Delia radicum* (HiC_scaffold 1 - 6) to homologues on the *Drosophila melanogaster* chromosomes (2R/2L, 3R/3L, 4, X and Y). Each line connects homologue regions of at least two consecutive genes. Colored lines indicate that homologue regions of a *D. melanogaster* chromosome are connected to those of the syntenic chromosome of *D. radicum*. Otherwise they are colored in grey.

In our synteny analysis, we successfully mapped the six pseudo-chromosomes of *D. radicum* to the six chromosomes of *D. melanogaster* (Figure 3b). This was achieved by predicting gene models (Table 1) in the *D. radicum* genome based on the annotated *D. melanogaster* genome using GeMoMa (Table S7). Genes annotated on the X chromosome of *D. melanogaster* mapped successfully on the second largest chromosome (HiC_scaffold_2) in the *D. radicum* assembly. Genes annotated for the other *D. melanogaster* chromosomes were mainly localized on the remaining four larger *D. radicum* chromosomes (Figure 3b). For the smallest chromosome (HiC_scaffold_6) we found indications that this might be related to chromosome 4 (NC_004353.4) of *D. melanogaster* (Table S7).

### 3.3 Genome annotation and functional gene annotation

#### 3.3.1 Process of genome annotation and evaluation

We sequenced the transcriptomes of all four life stages (eggs, larvae, pupae, and adults) of *D. radicum,* and included two stress factors (heat stress on adults and plant toxin on larvae) that are relevant for the survival of *D. radicum* to support the prediction of a comprehensive set of protein-coding genes in the *D. radicum* genome.

Our homology-based protein-coding gene prediction with GeMoMa relied on nine already sequenced and annotated genomes of phylogenetically related species, herbivore species sharing the same host plant range or common pests on crop plants or stored grains (Table 1). We predicted 19,343 protein-coding genes comprising 46,286 transcripts (Table 3) having a homologue in at least one of the nine selected species.

**Table 3.**
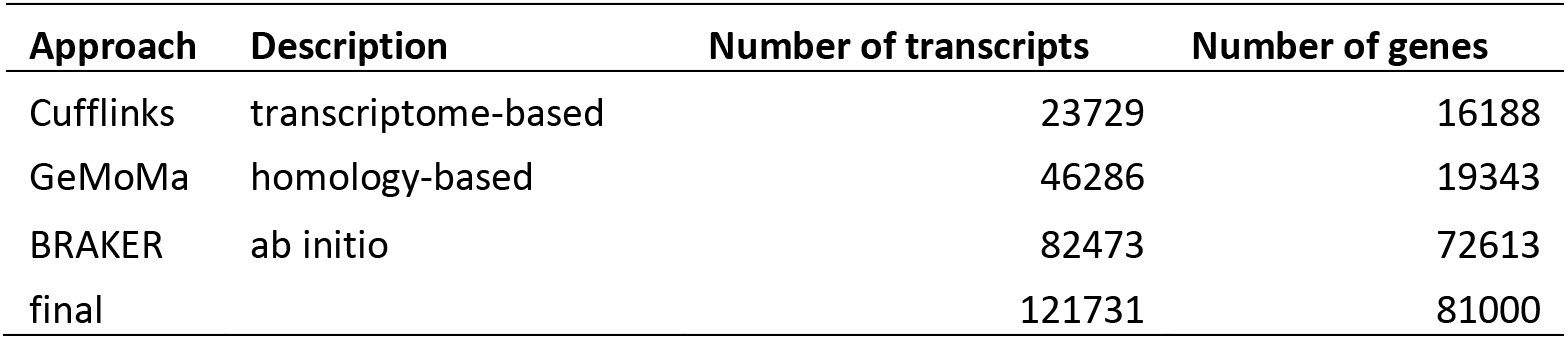
Summary of gene prediction statistics. Number of gene predictions made on the chromosome-scale genome assembly of *D. radicum* by the three different approaches: GeMoMa a sequence homology-based approach, Cufflinks a RNASeq data-based approach to assemble transcriptomes, and BRAKER an approach for *ab initio* predictions of genes. The final comprehensive gene annotation for the *D. radicum* genome contains 81,000 putative genes.

**Table 4.** Summary of Gene Ontology (GO) annotations of expressed genes. 30,492 genes of the 81,000 genes were expressed (TPM value ≥ 1) in our in-house life stage RNASeq data set. Genes were annotated with GO classes using PANNZER2 and InterProScan. Genes exclusively expressed in one specific life stage were grouped into gene sets named according to the life stage. For the larvae and the adult stage were control and stress conditions are present in the data set, genes that are expressed in both conditions within one life stage were added to the life stage and condition specific gene sets. Numbers of the row labeled with “total” are in concordance with Figure 5b. Listed gene sets were used for life stage specific GO enrichment analyses.

| set/ontology | complete | egg<br>(control) | larva<br>(control) | pupa<br>(control) | adult<br>(control) | larva<br>(ITC) | adult<br>(heatshock) |
| --- | --- | --- | --- | --- | --- | --- | --- |
| total | 30492 | 1457 | 2831 | 1770 | 1718 | 2895 | 2045 |
| noGO | 15222 | 1029 | 2021 | 1288 | 1096 | 2079 | 1225 |
| GO | 15270 | 428 | 810 | 482 | 622 | 816 | 820 |
| BP<sup>†</sup> | 11262 | 262 | 501 | 289 | 406 | 495 | 534 |
| MF<sup>‡</sup> | 13513 | 397 | 717 | 429 | 537 | 723 | 729 |
| CC<sup>§</sup> | 10917 | 230 | 444 | 260 | 381 | 434 | 454 |

<sup>†</sup> BP: Biological Process, <sup>‡</sup> MF: Molecular Function, <sup>§</sup> CC: Cellular Compartment

As a complementary approach, we assembled the transcriptomes of all life stages from egg to adult, plus adults and larvae subjected to two stage-related stress factors using Cufflinks. From the pure RNASeq-based transcriptome data, we were able to predict 16,188 protein-coding genes covering 23,729 transcripts (Table 3) that were expressed at the sampled time points of the different life stages.

To cover the hitherto non-annotated and not or low expressed *D. radicum*-specific genes under our conditions, we performed *ab initio* gene prediction. A total of 81,150 genes yielding 82,473 transcripts were predicted (Table 3). Similarly, as before, we retained all predicted genes, to allow future users the option to choose their own filtering criteria in later studies.

The integration of the predictions of all three approaches into a comprehensive annotation led to 81,000 putative genes covering 121,731 transcripts (Table 3), where a relatively high number of putative genes was predicted specifically by the *ab initio* approach (Figure 4a). Nearly 95.5 % of the genes were located on the six chromosomes (Table S3).

**Figure 4.**
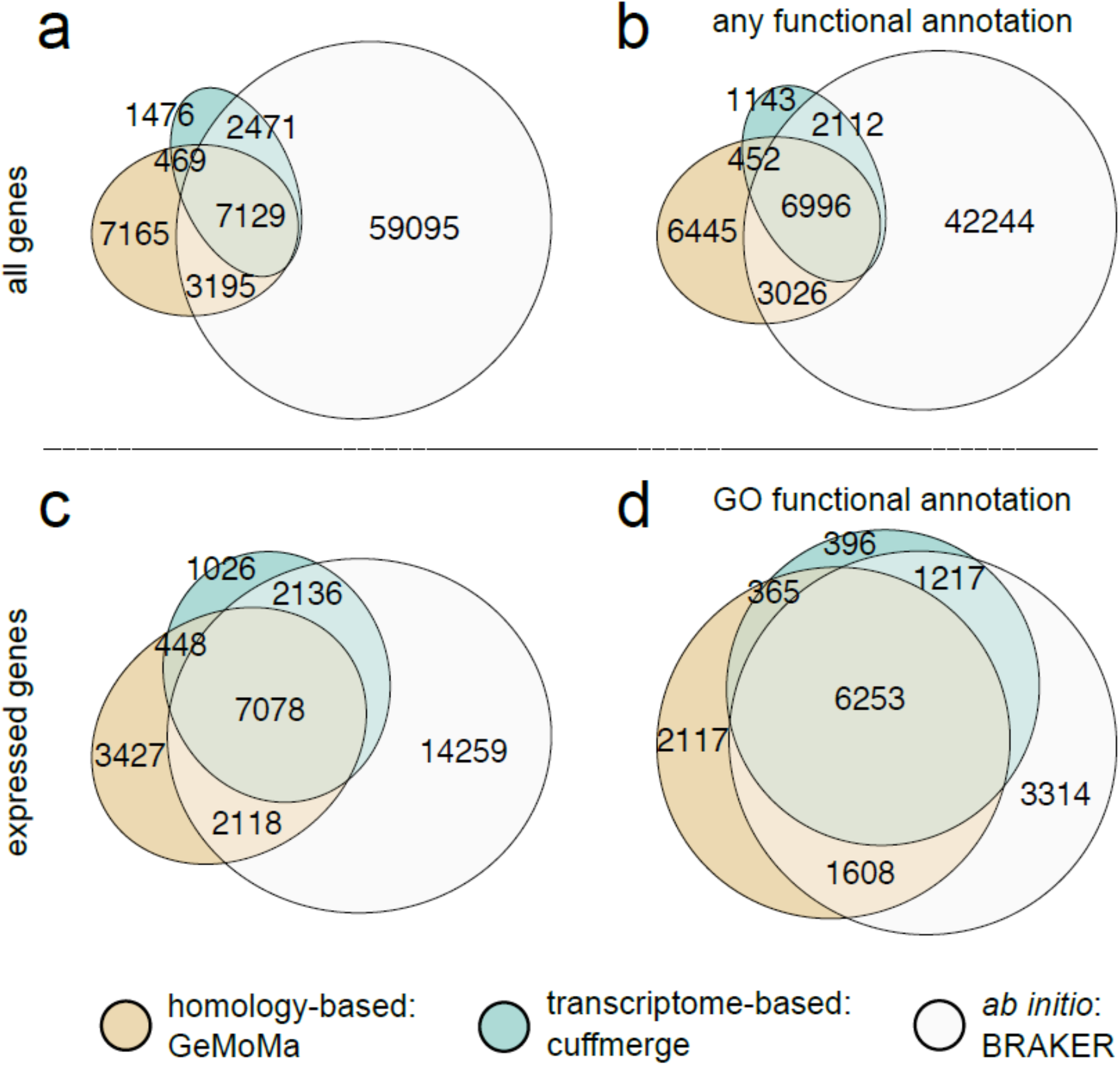
Venn diagrams containing the numbers of genes in the *Delia radicum* genome predicted by homology-based, transcriptome-based or *ab initio* approaches, or a combination thereof. The numbers of genes in the diagrams are based on (a) all predicted genes; (b) all predicted genes with any functional annotation, which includes GO annotation and/or protein family or domain annotation; (c) expressed genes which were predicted genes with a Transcript Per Million (TPM) value ≥ 1. TPM values result from analyses of our in-house life cycle RNASeq data.; (d) expressed genes with I think that is a great a functional annotation based on GO annotation.

Evaluation with BUSCO showed that our genome annotation covered 93.6 % complete-copy genes of the Diptera gene set, and 95.4 % of the Endopterygota gene set (Table S8). By determining the overlap of the predictions, we found 7,129 genes that were predicted by all three approaches and a total of 13,264 genes by at least two approaches (Figure 4a). The annotation of the latter set of genes covered 87.5 % of complete-copy genes of the Diptera gene set and 89.5 % of the Endopterygota gene set (Table S8). Only the combination of all three approaches led to a complete annotation of the *D. radicum* genome.

#### 3.3.1 Functional annotation

Overall, 77.1 % (62,418) of the genes were functionally annotated with at least one GO-term and/or protein family or domain information (Figure 4b, Figure S2, Table S9), including 71.15 % (42,244) of the only *ab initio* predicted genes.

Focusing on the expressed genes by using our in-house whole life stage RNASeq data, we found a reasonable number of 30,492 genes (37.64 %) having an estimated expression of ≥ 1 transcript per million (TPM) (Figure 4c). A high number of genes was predicted by BRAKER only, but most of these genes were not expressed under our conditions, although the number of expressed genes is higher than in the other sets (Figure 4c). From the set of expressed genes, 50 % (15,270) were functionally annotated with at least one GO-term (Figure 4d).

Taken together, these findings indicate that our gene annotation is complete and accurate. We will demonstrate its applicability to generate biologically relevant information in the following section, where we analyzed the transcriptomes of all life stages of *D.* radicum to identify life stage-specific functional gene expression underlying adaptations to their stage-specific life styles, especially to their host plants.

### 3.4 The *D. radicum* life cycle

An unsupervised clustering analysis of the expressed gene set with UMAP showed a high similarity of samples belonging to the same life stage (larva or adult, Figure 5a), even if they were subjected to different conditions (control and stressed). We also found that all samples of the egg and pupal stage clustered together. This seems logical, considering that the egg and pupal life stages both undergo considerable morphological and physiological transformation processes, and, in contrast to larvae or adults, are less involved with digestive, locomotory, gustatory and olfactory processes.

**Figure 5.**
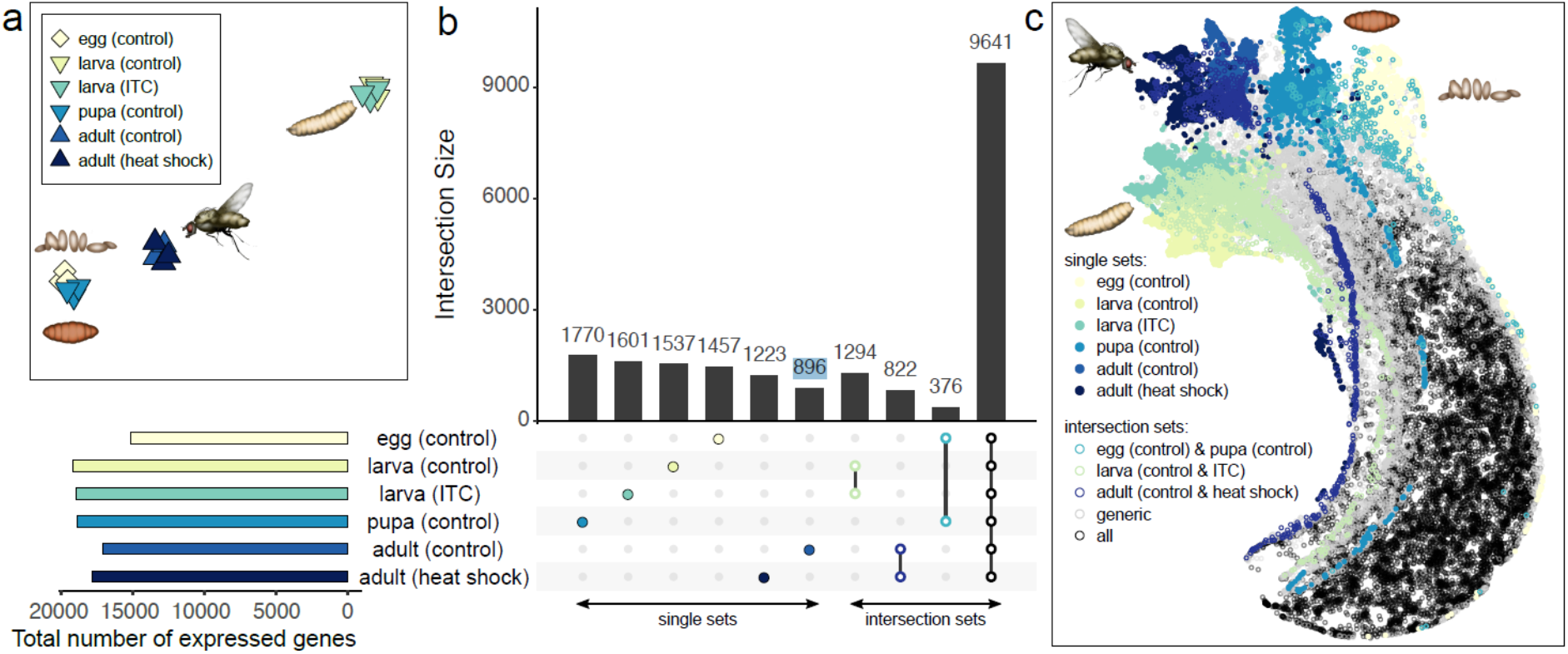
Differences in gene expression profiles among *Delia radicum* life stages and stress conditions. (a) Uniform Manifold Approximation and Projection (UMAP) plot showing differences among the life stages based on differing gene expression. ITC = larvae fed on diets with 2 μM phenylethyl isothiocyanate in their diet. (b) UpSet plot showing the number of genes that are exclusively expressed (Transcripts Per Million (TPM) value ≥ 1) in at least one replicate of a life stage or a stress condition (first 6 bars, filled circles); expressed in both conditions within larva and adult life stage (bar 7 and 8, green and dark blue open circles), or both in eggs and pupae (bar 9, cyan open circles), and those expressed in all 18 samples (last bar, open black circles). A selection of intersection sets is shown, whereas the full set is presented in Figure S4. To the left, the total number of expressed genes per life stage and stress condition is shown (colored horizontal bar plot below sub figure a). The remaining genes, referred to as generic, are not shown and sum up to 9,875 genes. (c) UMAP of expressed genes. Genes are colored according to the sets in (b) and are plotted with filled circles when they belong to single sets and with open circles when they belong to intersection sets. Genes expressed in all 18 samples are labeled as “all” (black open circles). Remaining genes are labeled as “generic” (grey open circles).

We also found that the total number of expressed genes differed among life stages (Figure 5b). The lowest total number of expressed genes was detected in the egg stage and the highest in the larval and pupal stages (Figure 5b, horizontal bar plot). When looking at the overlap among the life stages, we found 31.6 % of the 30,492 genes to be expressed across all life stages (Figure 5b, vertical bar plot). Another 36 % of the genes were exclusively expressed in either a single life stage or condition, in the intersection of both conditions of the larval and the adult stage, respectively or specific in egg and pupal stage (Figure 5b, Figure S3). In the UMAP plot (Figure 5c), genes expressed in single life stages were located at the top and formed life stage-specific spots, whereas genes expressed in all life stages also clustered but were located on the opposite side. The remaining one-third of the genes (not shown in Figure 5b, Figure S3) clustered in between. For larval or adult stages, we observed that genes expressed under different conditions clustered closely together and formed life stage-specific clusters (Figure 5c).

An ontology-based gene expression analysis revealed life-stage specific groups related to biological processes (BP), molecular functions (MF) and cellular components (CC, Figure 6, Figure S4, Table S10). In the egg stage, mainly genes involved in the embryonic development (BP), transcriptomic activity (MF) and genetic material (CC, Figure 6) were expressed. Especially genes belonging to the GO biosynthetic processes DNA biosynthesis, metabolic processes, egg shell layer formation (amnioserosa formation) and organ development (muscle and organ formation) were expressed (Figure S4a). These processes are involved in the transition from embryo to larva, which requires active cell division and involves a broad range of metabolic processes to synthesize cell components, membranes and organs (Beutel, Friedrich, Yang, & Ge, 2013). These structures require different macromolecules; indeed we found several expressed genes related to molecular biosynthesis processes in the eggs (Figure S4a). Cell differentiation and organ formation require regulation, coordination and binding activation (Izumi, Yano, Yamamoto, & Takahashi, 1994) which was reflected in our BP expression data (Figure 6, Figure S4a).

**Figure 6.**
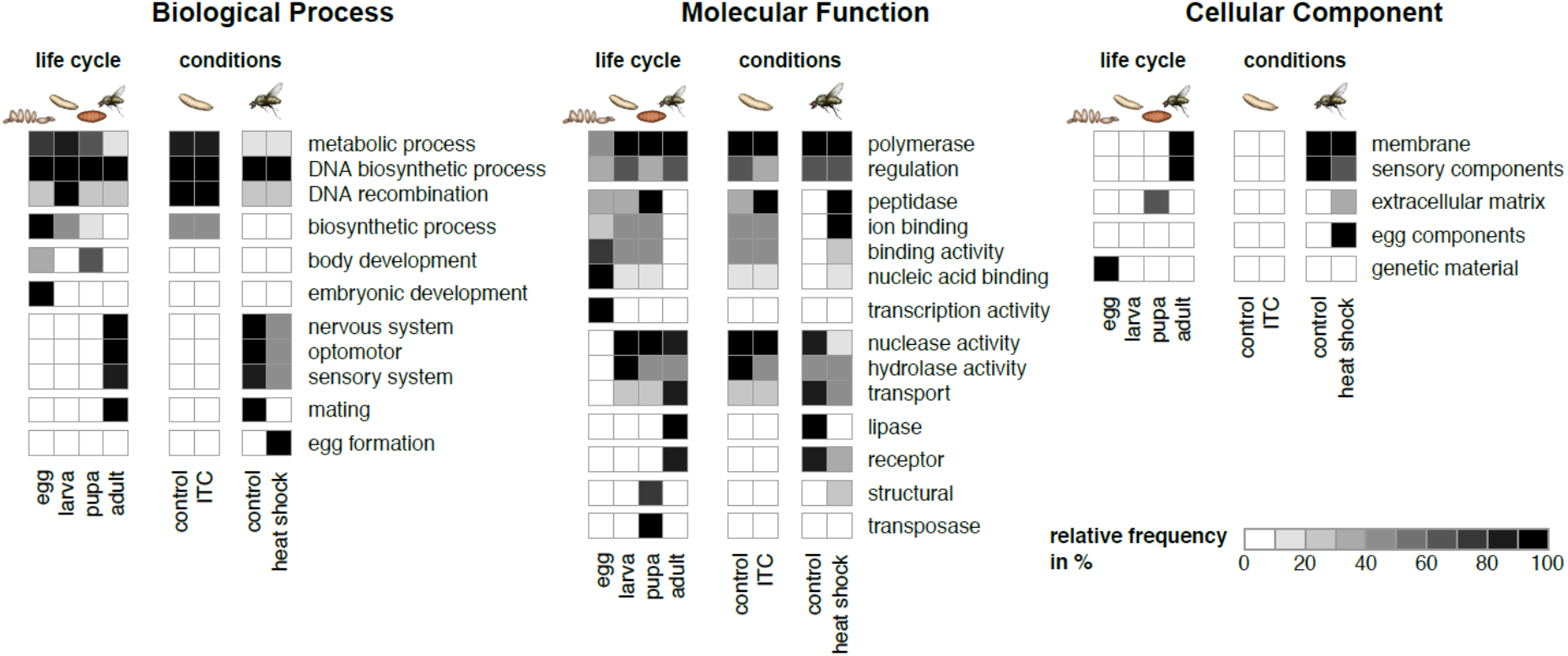
Gene ontology (GO) analyses on biological process (BP), molecular function (MF) and cellular component (CC) ontologies, based on expressed genes (Transcripts Per Million (TPM) value ≥ 1). Results are shown in three respective heatmaps, where rows are labeled by generic categories and columns with life stages and/or conditions. Explicit GO annotations of expressed genes are collapsed into more generic categories. Hence, each cell in a heatmap contains the relative frequency of GO terms sorted into a specific generic category for a specific life stage and/or condition. Only GO terms that were significantly overrepresented in a GO-enrichment analysis (Fisher’s exact test, P<0.05 after correction with Benjamini - Yekutieli) are considered. Expanded versions of the heatmaps, where detailed GO annotations for each generic category are listed, are provided in Figure S4. In all heatmaps the block with four columns to the left shows the results of all stage of the life cycle under control conditions, whereas the columns to the right show the relative frequencies determined for larvae and adults under control or stress conditions; the data for the control conditions in larva and adult stage are duplicated for easier comparison. ITC = larvae fed on diet with 2-phenylethyl isothiocyanate.

Genes involved in the body development (BP), structural and transposase (MF), and extracellular matrix (CC) were more frequently expressed in pupae (Figure 6). These genes belong to GOs comprising regulators, binding activity, biosynthesis, metabolism and DNA amplification (Figure S4). During the pupal stage, metamorphosis results in the ‘disassembly’ of larval structures to form adult wings, compound eyes and legs (Buszczak & Segraves, 2000; Chapman & Chapman, 1998). This requires the expression of genes involved in catabolic processes, as well as in organ and cuticle formation. Indeed, we found an increased expression of genes responsible for nuclease and peptidase activity (MF) and chitin-based cuticle structures (CC, Figure 6, S4). This is in line with the gene expression profiles in *D. melanogaster* pupae (Arbeitman et al., 2002).

Genes connected to the metabolic processes (BP) were highly expressed in the larval stage (Figure 6). We found genes coding for peptidases and polymerases (MF), involved in DNA processes or functions and biosynthetic processes (BF) to be highly expressed (Figure 6, Figure S4). These genes are likely related to feeding and digestion as well as to growth and molting, which are the main processes in the larval stage (Chapman & Chapman, 1998; Chen, 1966). In larvae exposed to the plant toxin ITC, we found that peptidase genes (MF) and genes involved in metabolic and biosynthetic processes (BP) were activated (Figure 6, Figure S4). These enzymes may be involved in catabolizing plant toxins as has been described for other herbivores feeding on *Brassica* plants (Schramm, Vassão, Reichelt, Gershenzon, & Wittstock, 2012).

Genes coding for the detection of visible and UV-light, optomotor capability, detection of chemical stimuli (taste, smell) and temperature (BP) were exclusively expressed in adults (Figure 6, Figure S4a). The expression of these gene sets, which are involved in the sensory, optomotor and nervous systems (BP), are important to localize food sources and suitable hosts for oviposition (Gouinguene & Städler, 2005; Gouinguené & Städler, 2006; Peter Roessingh & Städler, 1990). In addition, several genes coding for receptors and ion channels were expressed (Figure 6, MF). These genes are involved in the detection of environmental stimuli and signal transmission via the nervous cells to the brain (Sato & Touhara, 2008). Specific for the adult life stage were also the expression of adult behavior linked genes (Figure S4a).

Exposing adult flies to a higher temperature resulted in the enhanced expression of peptidases, ion binding (MF), sensory system, especially smell and egg formation (BP) related genes compared to control adults (Figure 6, Figure S4). High temperatures alter protein stability, structures and folding, followed by functional changes (Jaenicke et al., 1990). The activation of peptidases might avoid malfunction of proteins under heat stress. Temperature changes affect also the volatility of volatile organic compounds (VOCs) as well as the emission rates of plants (Copolovici & Niinemets, 2016). Since adults of *D. radicum* are attracted by VOCs to localize host plants (Finch, 1978), the enhanced expression of genes related to VOC perception (smell) in flies might indicate towards an adaptive response. Investing in offspring, under these circumstances might ensure the survival for the fly population, and to localize a possible host plant for their oviposition, *D. radicum* females utilize odor signals (Nottingham, 1988).

## 4 Conclusion

An increasing number of assembled and annotated insect genomes have been published over the last decade. However, genomes of belowground insects and especially root-feeding herbivores are underrepresented. We sequenced the genome of a belowground-feeding agricultural pest, the cabbage root fly *Delia radicum,* whose larvae are also used as a ‘model’ belowground herbivore in studies on optimal defense allocation and systemic induced responses in plants. Using PacBio and Hi-C sequencing, we generated a 1.3 Gbp assembly with an N50 of 242 Mbp, 6 pseudo-chromosomes and 13,618 annotated genes using homology-, transcriptome- and model-predicted approaches, predicted by at least two approaches. During the assembly process, we identified a co-assembled *Wolbachia* species, a very common endosymbiont in insects (Werren & Windsor, 2000). The *Wolbachia* genome consisted of a single contig of 1.59 Mbp matching to the size of the *Wolbachia* supergroups A and B (~1.4 − 1.6 Mbp) which are typical for arthropods (Lo, Casiraghi, Salati, Bazzocchi, & Bandi, 2002). Such co-assembled endosymbiont genomes can be valuable to understand host-symbiont interactions and their roles in other interactions such as host-plant adaptations.

Our accurate and well-annotated genome can reveal genetic mechanisms underpinning adaptation of *radicum* to its host plants and their specific chemical defenses, the glucosinolate-isothiocyanate system. With our work we provide a tool to understand how the different life stages of this herbivore have adapted to their host plants by identifying adult-specific genes involved in olfactory orientation or the detoxification of plant defense compounds in larvae. The genome and the transcriptomes can further be used to understand adaptation to specific conditions, i.e. the evolution of pesticide resistance and adaptive responses to environmental stress factors, such as temperature increase or soil pollution. This high-quality genome is also an important tool to develop novel strategies to combat this pest, for example highly specific dsRNA-based pesticides, which can discriminate between target and non-target species. Moreover, the genus *Delia* contains several other pest species, such as the turnip root fly *D. floralis*, the onion fly *D. antiqua* and the seed bulb maggot, *D. platura.* As their common names indicate, they attack a range of agricultural crops. The genome of *D. radicum* is an excellent foundation to further explore the genetic mechanisms underlying adaptation to chemical host-plant defenses among member of the genus *Delia*.

## Acknowledgements

We thank the Long Read Team of the DRESDEN-concept Genome Center, DFG NGS Competence Center, part of the Center for Molecular and Cellular Bioengineering (CMCB), Technische Universität Dresden and the MPI-CBG, especially Sylke Winkler for their great support and the collaboration. Dominik Jakob is acknowledged for his assistance with the insect culture. Great thanks to Denis Tagu and Fabrice Legeai (INRAE, Rennes, France), Denis Poinsot (University of Rennes, France) and Ekaterina Shelest (BIU, iDiv, Leipzig, Germany) for their helpful advice and encouragement in earlier stages of this project. We also greatly thank Jens Keilwagen (JKI, Quedlinburg) for valuable discussions on gene prediction and support on GeMoMa.

## Author Contributions

YP, RS, ND designed the project, AC provided the starting culture of the insects, RS, HV performed the laboratory work, YP, CG, YO, HV performed the data processing and analysis, YP created the figures, YP, RS, ND, HV wrote a first version. All authors contributed to the writing process.

## Data Availability Statement

Genome sequences and raw data used for genome assembly (PacBio sequences and Illumina Hi-C sequences) and annotation (Illlumina RNASeq sequences) will be available at NCBI. Final genome annotation, respective annotations by GeMoMa, Cufflinks and BRAKER, and functional transcript annotations made by InterProScan and PANNZER2 will be available via Zenodo.

