## Supplementary material for "A high-quality functional genome assembly of *Delia radicum* L. (Diptera: Anthomiidae) annotated from egg to adult": Figure S1

DNA extraction from *D. radicum* for PacBio sequencing

Fly tissue was added to a lysis buffer containing 400 nm NaCl, 20 mM Tris HCl (pH 8.0), 30 mM EDTA (pH 8.0) and carefully mixed. Afterwards, 125 µl SDS (10%) and 15 µl Proteinase K (10 mg/ml) were added to the mixture and incubated over night at 55°C. The next day, samples were centrifuged for 30 min at 4000 x g at room temperature (RT) and the supernatant transferred to a new tube. We added 5 µl RNAse A (2 mg/ml) to the supernatant, mixed carefully and incubated the samples for 1 h at 37°C. After this step, the lysed tissue was washed twice with 250 µl Phenol:Chloroform:Isoamylalcohol (25:24:1, equilibrated with 10 mM Tris, pH 8.0 and 1 mM EDTA) and carefully mixed on an Intelligent Mixer for 10 min at RT. In the next step, the samples were centrifuged for 10 min at 4,000 x g at RT and the aqueous phase transferred to a new tube. The aqueous solution was washed with 250 µl Chloroform, carefully mixed and centrifuged at 4,000 x g, at RT for 10 min. The last washing step was repeated twice. Afterwards, the gDNA was precipitated by adding 450 µl precooled 96% ethanol, carefully mixed and centrifuged at 4,000 x g at 6°C for 20 min. The supernatant was discarded and the pellet washed twice with 400 µl 70% cold ethanol. Samples were shortly spun, dried for 10 min at 37°C and dissolved in 50 µl 1x TE-buffer.
