## Supplementary figures and images for "A high-quality functional genome assembly of *Delia radicum* L. (Diptera: Anthomiidae) annotated from egg to adult"

### Figure S2

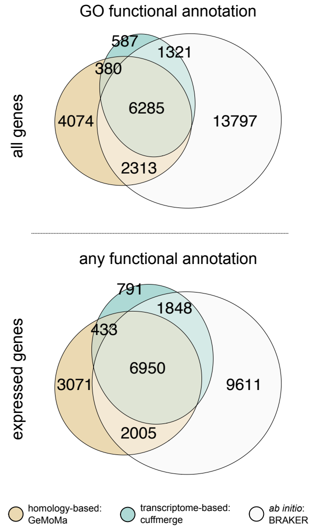

### Figure S3

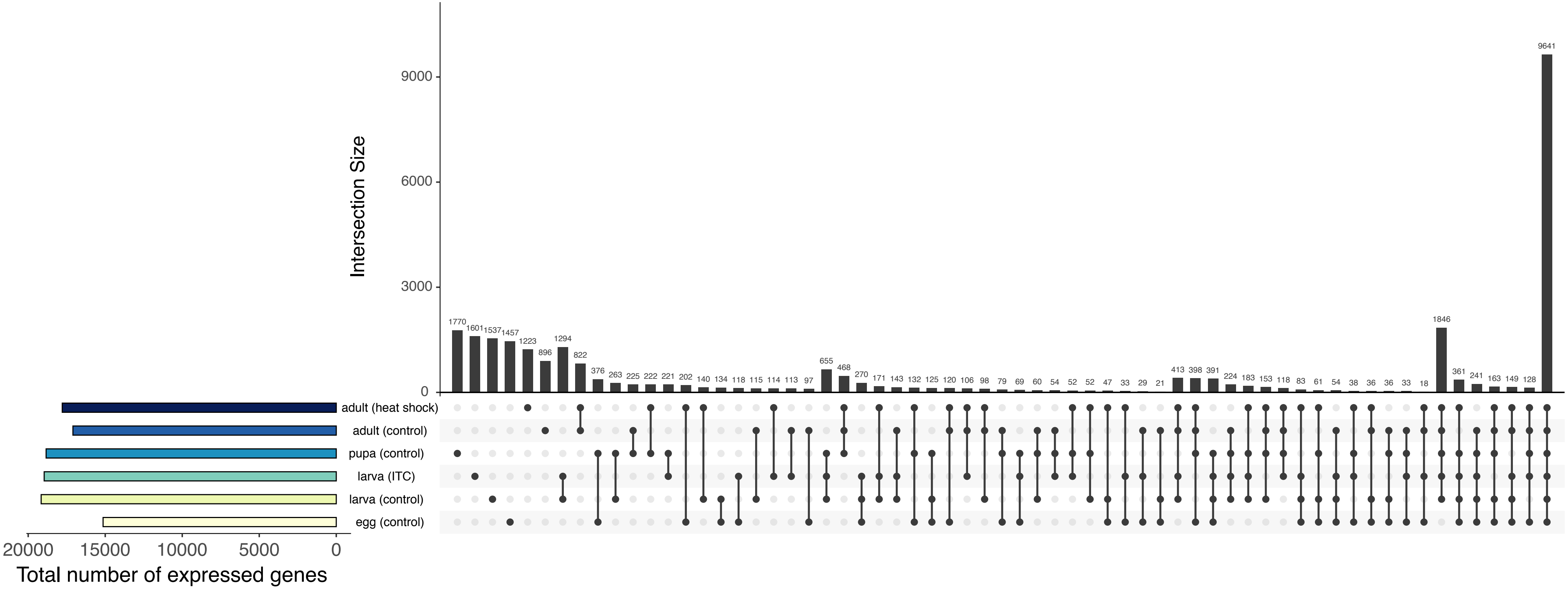

### Figure S4

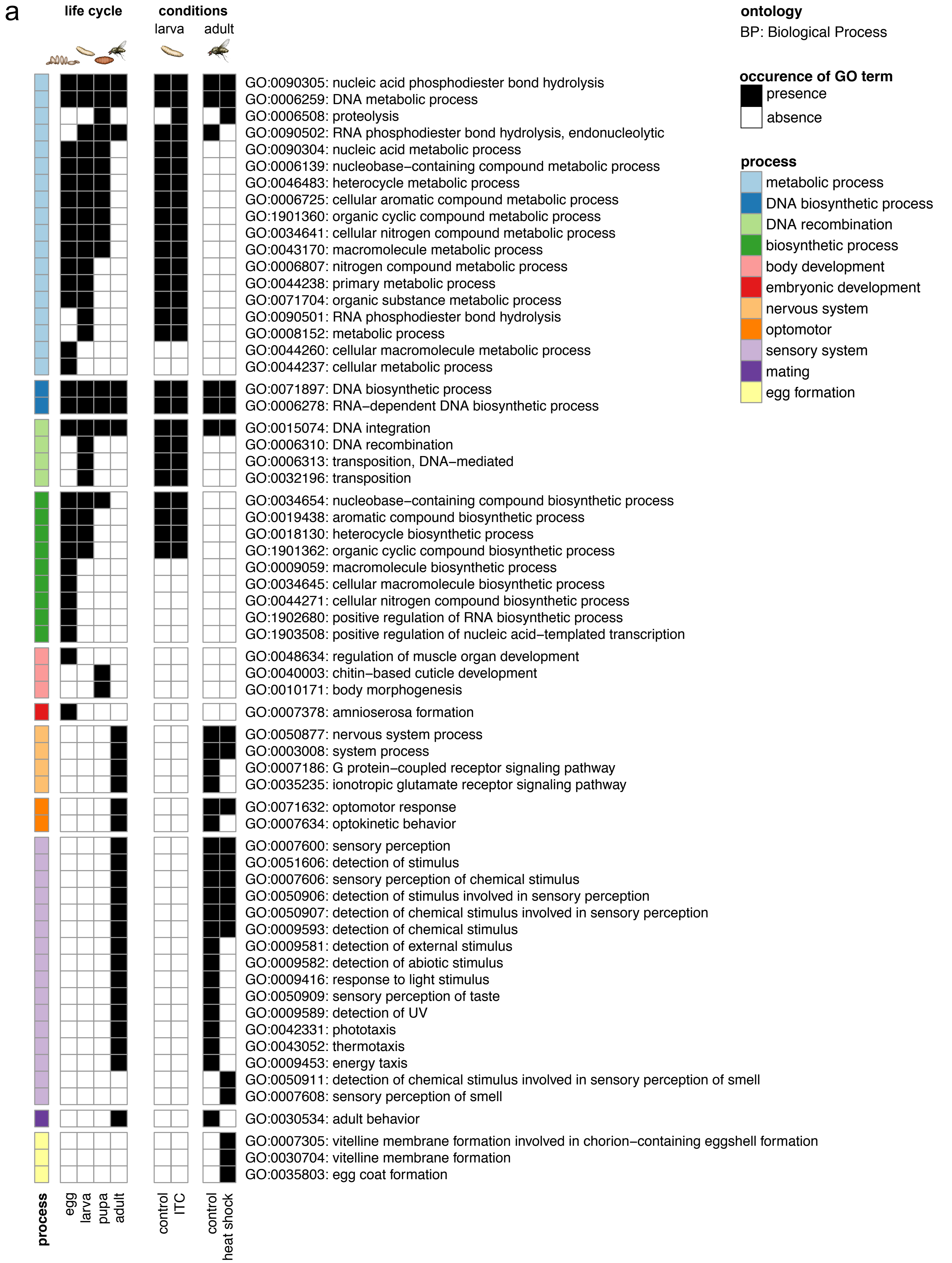

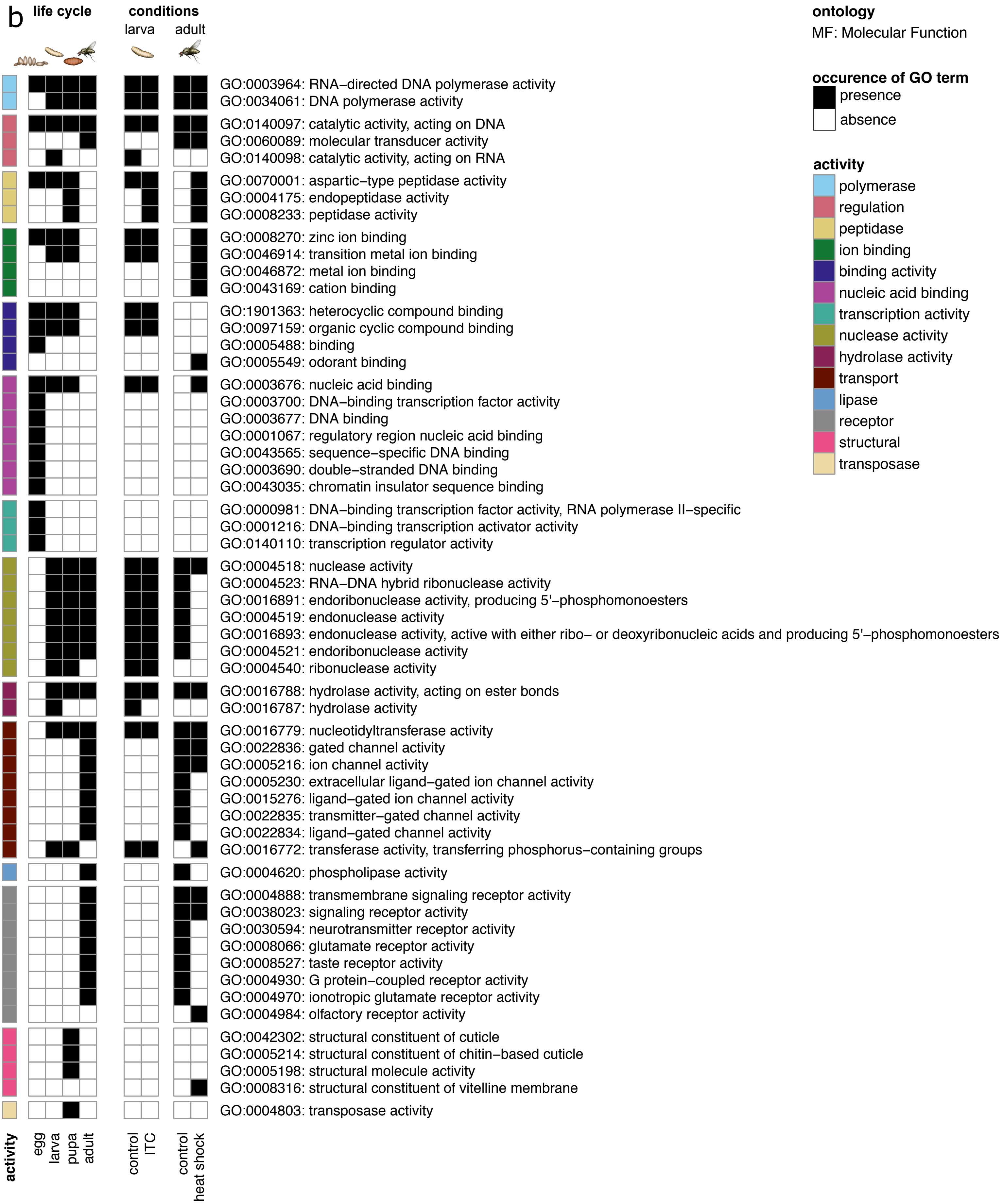

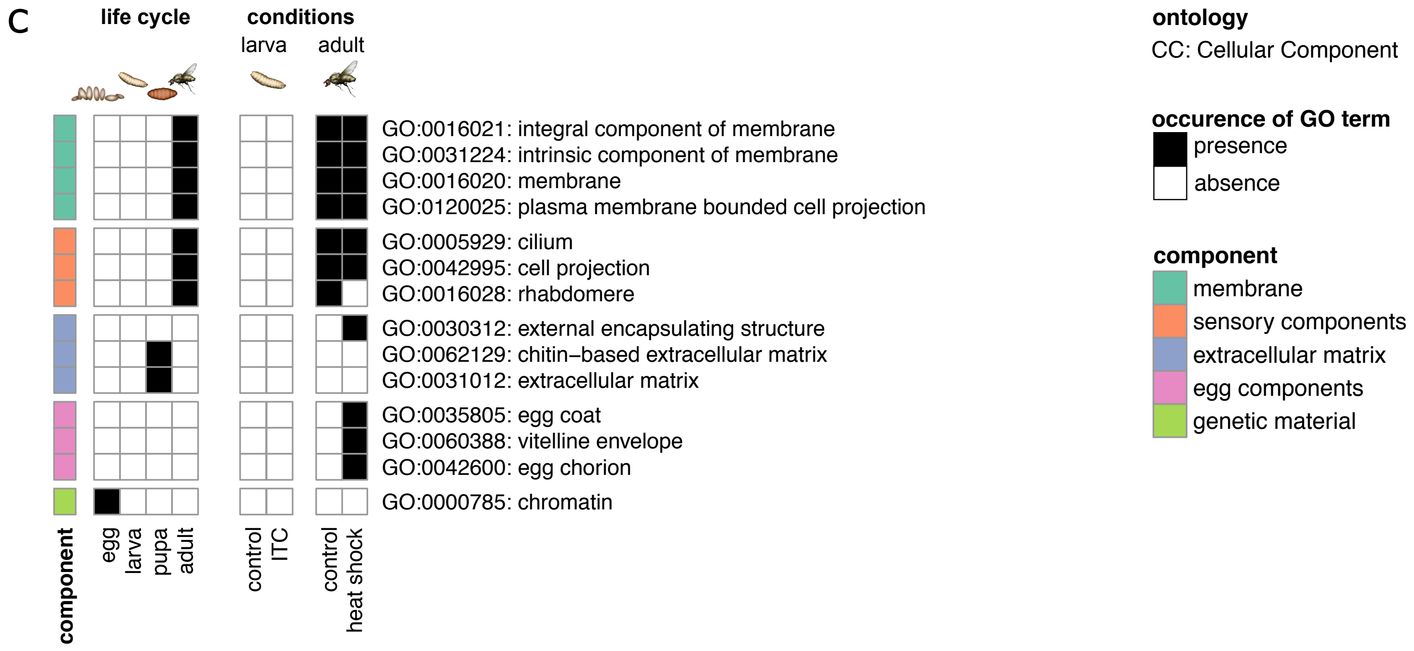
